# TEsingle enables locus-specific transposable element expression analysis at single-cell resolution

**DOI:** 10.64898/2026.03.19.712984

**Authors:** Talitha Forcier, Esther Cheng, Oliver H Tam, Cole Wunderlich, Laura Castilla-Vallmanya, Joanne L. Jones, Annelies Quaegebeur, Roger A. Barker, Johan Jakobsson, Molly Gale Hammell

## Abstract

Transposable elements (TEs) are mobile genetic sequences that can generate new copies of themselves via insertional mutations. These viral-like sequences comprise nearly half the human genome and are present in most genome wide sequencing assays. While only a small fraction of genomic TEs have retained their ability to transpose, TE sequences are often transcribed from their own promoters or as part of larger gene transcripts. Accurately assessing TE expression from each individual genomic TE locus remains an open problem in the field, due to the highly repetitive nature of these multi-copy sequences. These issues are compounded in single-cell and single-nucleus transcriptome experiments, where additional complications arise due to sparse read coverage and unprocessed mRNA introns. Here we present our tool for single-cell TE and gene expression analysis, TEsingle. Using synthetic datasets, we show the problems that arise when not properly accounting for intron retention events, failing to address uncertainty in alignment scoring, and failing to make use of unique molecular identifiers for transcript resolution. Addressing these challenges has enabled an accurate TE analysis suite that simultaneously tracks gene expression as well as locus-specific resolution of expressed TEs. We showcase the performance of TEsingle using single-nucleus profiles from substantia nigra (SN) tissues of Parkinson’s Disease (PD) patients. We find examples of young and intact TEs that mark dopaminergic neurons (DA) as well as many young TEs from the LINE and ERV families that are elevated in PD neurons and glia. These results demonstrate that TE expression is highly cell-type and cellular-state specific and elevated in particular subsets of neurons, astrocytes, and microglia from PD patients.

## Introduction

Transposable elements (TEs) are mobile genetic elements that comprise at least 45% of the human genome^1^. These viral-like sequences lay dormant within most healthy somatic cells but have been shown to be transcriptionally active in aging tissues^2,3^ and in the context of disease like cancer^4–7^ and neurodegeneration^8–17^. Tracking TE activity across tissues and disease has been stymied by a lack of rigorous tools for accurately assessing expression from these highly repetitive genomic elements^18,19^. Recently, several groups have developed tools to address the difficulties associated with properly handling TE reads. This includes software packages for tracking TE reads in bulk^20–36^ and single-cell^37–40^ RNA-seq data (scRNA-seq). However, accurate and sensitive detection of TEs in single-cell genomics datasets remains an ongoing problem.

TEs in single-cell data remain challenging to quantify due to their highly repetitive sequences and presence as multiple copies across the genome, with approximately 25% of TE copies located within introns of expressed genes^41^. These characteristics make it difficult to accurately assign TE-derived reads to the correct locus of origin and to distinguish true TE expression from intron retention events or unprocessed pre-mRNA. Accurate handling of TE reads becomes especially important when trying to analyze data from single-nucleus RNA sequencing (snRNA-seq) assays where unprocessed intron sequences constitute up to 40% of the transcripts captured – and these intronic sequences are packed with both functional and fragmentary TEs. Given these challenges, accurate quantification of TE-derived transcripts for scRNA-seq data requires computational tools that can accurately align multi-mapping reads, perform UMI correction while accounting for ambiguity in alignment and annotation, properly handle intron inclusion, and accurately assign ambiguous reads to their locus of origin.

Here we present a new software package for the accurate assessment of TE expression in scRNA-seq and snRNA-seq. We additionally developed methods for generating realistic single-cell and single-nucleus simulated data to benchmark the performance of TEsingle. We show that TEsingle simultaneously assesses both gene and TE expression and optimizes for TE analysis without compromising its accuracy on gene abundance estimation. In fact, TEsingle outperforms other standard single-cell analysis software by also returning better estimates for gene expression. We next applied TEsingle to a publicly available postmortem snRNA-seq dataset from the substantia nigra pars compacta (SNpc) of Parkinson’s Disease patients to showcase the ability of TEsingle to recover TE expression associated with PD. Results from this analysis demonstrated that TE expression is highly cell-type specific and elevated in specific subpopulations of neurons, astrocytes, and microglia in PD patient tissues, signaling a potential role for TEs in PD progression.

## Results

### 1. The TEsingle algorithm

TEsingle focuses on accurate assignment of aligned reads to the most likely gene or TE locus of origin. The basic design of TEsingle follows our popular software package for bulk gene expression analysis, TEtranscripts^20^. More specifically, the heart of the TEsingle algorithm involves an expectation-maximization algorithm for resolving any ambiguous alignments of TE-derived RNA-seq reads. This method was originally developed for TEtranscripts and has since been adopted for TE expression analysis by many groups in the field^28,34,35^. To adapt TEtranscripts for single-cell and single-nucleus RNA-seq data, we added several steps to properly handle Unique Molecular Identifier (UMI) barcodes that label individual captured transcripts and enable resolution of PCR duplicates in snRNA-seq data. UMIs are randomized nucleic acid barcodes built into the sequencing primers to help identify all reads produced from the same originating transcript^42,43^. Furthermore, we adjusted the output format to conform to Matrix Market Exchange (MEX) formats, enabling easy integration into downstream single-cell analysis packages. An overview of the workflow for the TEsingle algorithm is given in **Figure 1**.

**Figure 1:**
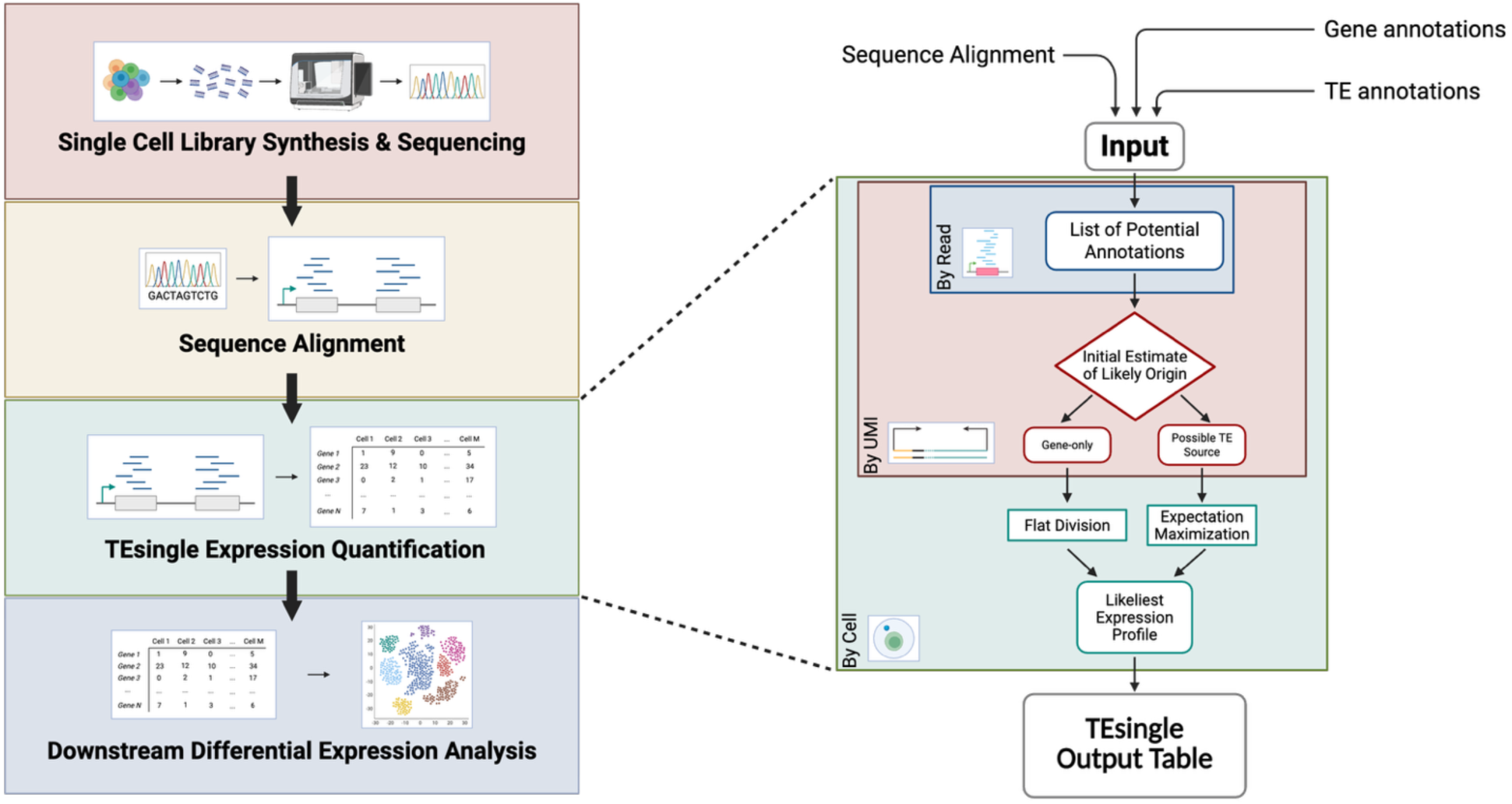
TEsingle Flow Chart. A flow chart depicting both the overall pipeline for scRNA-seq analysis and how TEsingle fits within it, with details of TEsingle’s internal processes. Genome aligned single-cell libraries using STARsolo and GTF files specifying locations of genes and TEs in the genome are used as inputs for TEsingle. After running TEsingle, a count table for UMI counts for each cell evaluated is outputted, which can then be further evaluated for differential expression. Mapped reads are organized first by cell barcode, second by transcript UMI, third by read name, and finally by alignment. Each alignment is first assigned a list of potential annotations and evaluated to determine if the transcript of origin could only have had a gene annotation source. Transcripts classed as gene-only are then assigned count values by flat division across potential annotations, weighted by number of reads and alignments giving evidence for each annotation. Transcripts classed as potentially of TE origin are then sent to an iterative algorithm to optimally disambiguate their annotation of origin. These count distributions are then evaluated to arrive at one annotation of origin for each transcript. Summed counts of all transcripts are then output for all cells.

As input, TEsingle requires an alignment file, in either SAM or BAM format^44^, and two annotation files, in GTF format^45^, to identify all annotated genes and TEs, respectively. The alignment file generated by STARsolo^46^ provides the algorithm with a whitelist-corrected set of cell barcodes and UMIs for each read alignment. With these input files, TEsingle first builds one index tree structure for gene annotations and locations, and a separate index for TE annotations and locations. Gene annotations are modeled as full- length transcripts, rather than excluding reads that align to intronic regions, which we find improves disambiguation of intron retention events from intronic TE expression.

TEsingle uses the whitelist-corrected cell barcode and uncorrected UMI flags to group reads by transcript. It then uses the searchable index structures to enumerate all possible annotations to which each read could have aligned. At this point, TEsingle then divides the sequencing data by individual cell barcode for further processing on a cell-by-cell basis. For each cell, TEsingle builds a UMI graph network, where each UMI is represented by a single node and edges connect any two nodes whose sequence differs by one Hamming distance. This enables TEsingle to group together any reads that likely derived from the same originating transcript, but which contain up to 1 PCR or sequencing error within the UMI sequence. Every independent subgraph is taken as one transcript, and the associated reads, for all the interconnected UMIs that make up that subgraph, are pooled to determine annotation of origin. From this point forward, TEsingle works at the UMI/transcript level, where each pooled UMI subgraph is assumed to represent a single originating transcript.

To determine the initial estimated annotation of origin for a given transcript, TEsingle first checks the pool of reads for that transcript’s UMI subgraph to find any reads that map only once. These uniquely mapping reads are then used to anchor the list of all possible annotations for that UMI/transcript group. For any UMI with at least one uniquely mapping read, the annotations supported by those unique reads define a restricted list of possible origins from either gene or TE sources. All reads associated with that UMI—both unique and multi-mapped—are then used to count how many are associated with each of the annotations on the restricted list. Then the UMI counts are normalized and proportionally distributed based on the fraction of reads that overlap each annotation. If a UMI has no uniquely mapping reads, then all of its reads are distributed over every annotation that it mapped to as an initial estimate. Each read for that UMI will contribute proportionally to the annotation it could come from, and after allocating these fractional contributions across all reads and all their possible annotations, the totals are normalized so the UMI contributes one full count.

With the initial estimated genomic origins established, each UMI with a nonzero value for any TE annotation is further processed using an expectation-maximization (EM) algorithm^47^ to determine the most likely annotation for each transcript. Transcripts with at least one uniquely mapping read assigned exclusively to a gene annotation are excluded from the EM algorithm to prevent potential biasing of TE-originating transcript counts towards abundantly expressed genes.

During development of TEtranscripts and TEsingle, we noted that inclusion of uniquely mapping gene and TE reads within the EM algorithm tends to incorrectly bias the outcome toward assignment of reads to genes (which tend to be more abundantly expressed) and older TE copies (which tend to contain more unique sequences)^20^. Thus, only ambiguous reads with potential TE annotations are included in the EM algorithm, a strategy that mirrors TEtranscripts. Once the EM algorithm finishes determining the transcript of origin for the pool of ambiguously mapped and annotated UMIs, the total counts for all UMIs are summed for each cell and arranged as an output table in cell by gene format, where “gene” includes all expressed genes and TEs detected.

### 2. Generating Simulated data for testing single cell analysis packages

We next set about constructing realistic synthetic datasets for testing the TEsingle algorithm (**Figure 2**) that would fully recapitulate the cell barcodes, UMIs, and gene sequence expected for the raw scRNA-seq and snRNA-seq data (**Figure 2A**). To ensure these simulations were faithful to real data, we evaluated the expression characteristics from publicly available sets of snRNA-seq data^48^ and whole-cell scRNA-seq data^49^. Our focus was on the relative expression levels, intron retention rates, and error rates within gene and TE mapping reads. We developed custom methods for calibrating absolute intron retention rates in various types of single-cell data (see methods) and determined the average intron retention rates to be 22.6% for whole-cell single-cell data and 41.4% for whole-nucleus (snRNA-seq) data (**Figure 2B**). See **Supplemental Table S1** for individual sample intron retention rates. Given the stark differences between the two data types, and the importance of disambiguating intronic TE expression from retained intron sequences, we decided to generate one set of synthetic data that used an intron retention rate of 20% for whole-cell scRNA-seq and a second set using an intron retention rate of 40% for whole-nucleus snRNA-seq simulations.

**Figure 2:**
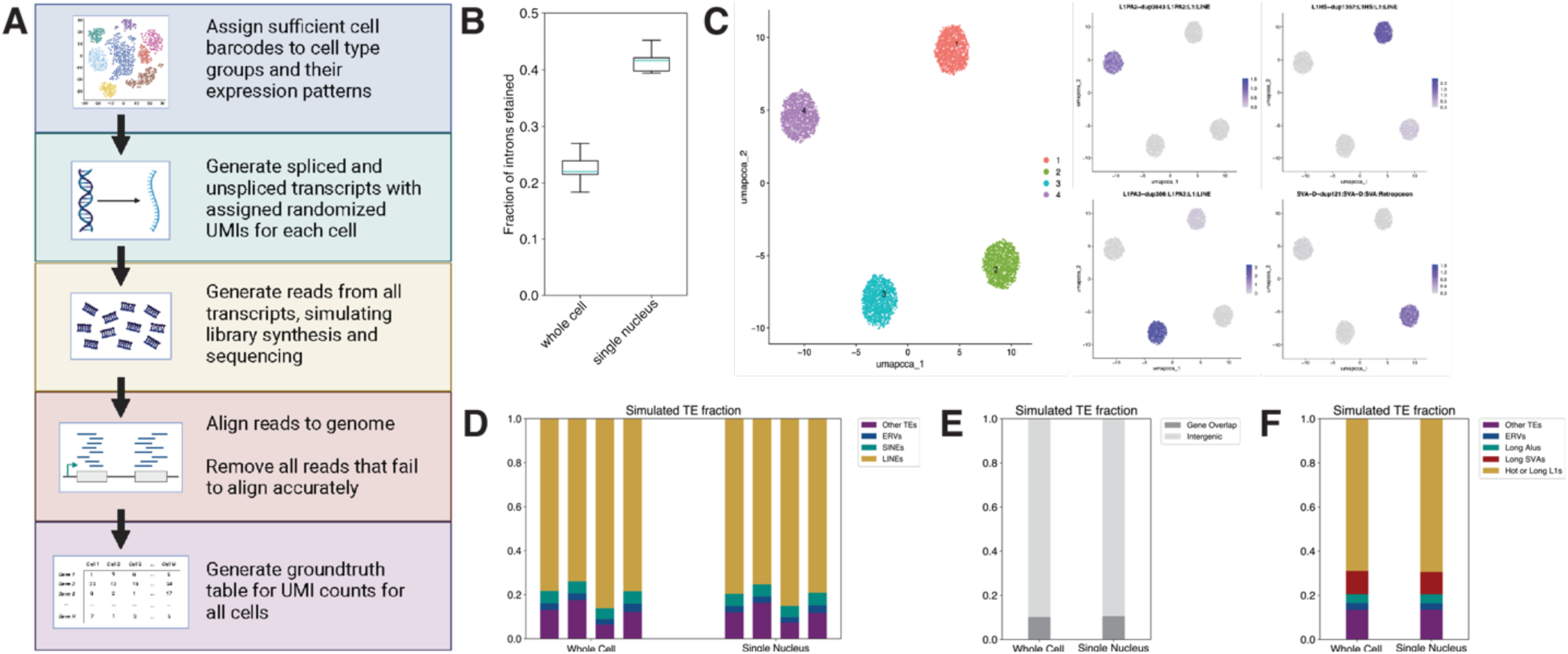
Simulated sequencing data characterization. **A:** A flow chart on how the simulated data was generated. **B:** Measured fractions of intron retention for publicly available scRNA-seq and snRNA-seq data. **C:** UMAP plots for all cell populations for the ground truth expression profiles of simulated snRNA-seq data. Additional UMAP plots included to showcase the four chosen TE loci expression profiles. **D:** Breakdowns of the number of TE transcripts with simulated expression in both simulated datasets. Whole-cell and snRNA-seq experiments were divided into four different cell type expression profiles, with separate simulations done for each type, giving different individual locus expression profiles for TEs but similar expression profiles for major TE classes. **E:** Breakdown of the fraction of TE transcripts simulated from either intergenic (light grey) or intronic (gene overlapping, dark grey) genome regions for both whole cell and single-nucleus simulated datasets. The majority of the simulated expressed TEs were from intergenic locations. **F:** Breakdown of the fraction of TE transcripts simulated from intact (promoter-containing) TEs of different types, including intact and full length: LINE-1s (yellow), SVAs (red), Alu/SINEs (green), ERVs (blue), and all other TE types (purple).

### 3. Benchmarking across TE expression analysis software

The simulated datasets described above were next used to determine the relative accuracy (precision and recall) in gene and TE expression estimates across a range of software packages capable of handling TE-derived sequences. To assess TEsingle’s performance in improving TE expression estimation, we compared it with STARsolo^46,50^ (version 2.7.11b), CellRanger^51^ (version 8.0.1), scTE^52^ (version 1.0), and soloTE^53^ (version 1.09). While some of these software packages were not specifically designed to handle TE reads by default (STARsolo and CellRanger), we included these packages if it was practical to alter the software packages to enable TE expression estimation without altering the core algorithms of that package. Each of these software packages differs in the strategies used for: aligning reads, handling ambiguously aligned reads, implementing genomic annotations, and resolving TE/gene annotation conflicts (**Table 1**). Furthermore, only two of these packages evaluate TE expression at the individual locus level for all TEs (**Table 1**). Further details on implementation of each package are given in the methods section.

**Table 1:**
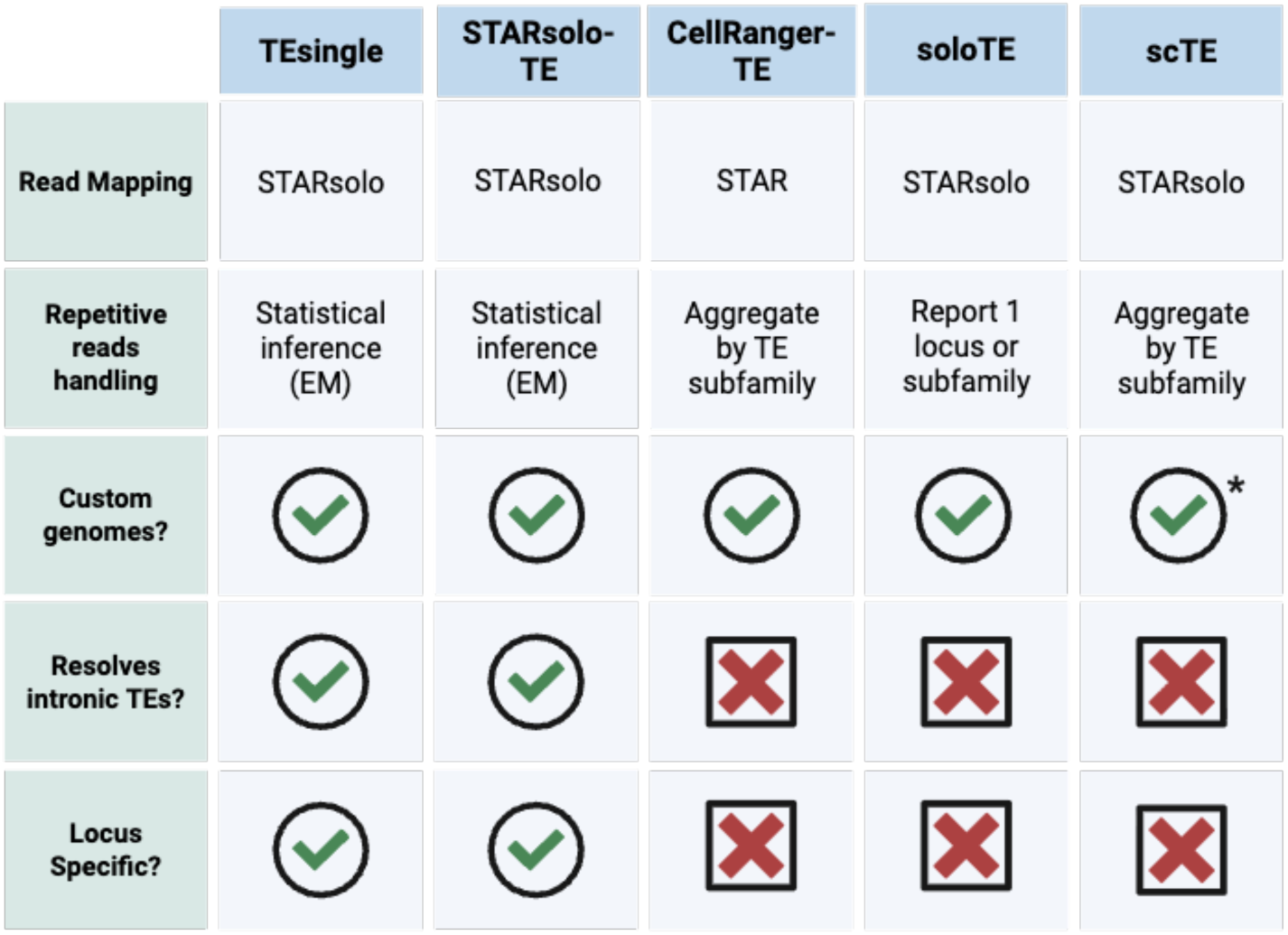
Software feature comparison. Available features for benchmarked software. STARsolo and CellRanger require tailored modified files to evaluate TE expression and were given the ‘-TE’ suffix to denote these changes. All software made use of STAR or STARsolo for initial alignment. How each software handled multimapping or repetitive reads was taken from the initial publication for each software. While soloTE can give some locus-specific counts for reads with sufficiently high mapping quality, its standard output contains a mixture of subfamily-level and locus-level counts.

To assess “accurate” expression estimates, we counted as accurate any gene or TE locus reported by each method whose estimated counts were within ±15% of the ground truth expression for that gene/TE locus. Any gene/TE with attributed counts that were greater than 115% of the simulated ground truth count were classified as ‘overestimated’ and those with counts less than 85% of the simulated ground truth value were classified as ‘underestimated’. Any gene/TE locus that was reported as expressed, but which was not part of the simulated truth set was classified as a ‘false positive’.

Conversely, loci that did have counts in the ground truth table but were predicted by any method to have 0 counts, were classified as a ‘false negative’ for that method. The breakdown of accuracy estimates for each benchmarked software package can be found in **Supplemental Figure S1C** for gene expression, **S1D** for TE expression when assessed at the TE subfamily level, and **S1E** for TE expression counted by individual TE loci. Relative rates for each category across all software tested are also shown in **Supplemental Tables S4, S5, and S6** for genes, TEs as measured by subfamily and TEs as measured by locus, respectively.

Using these categories, we measured accuracy for all gene/TE expression analysis packages by calculating the F1 score^54^ which considers both precision and recall in a manner that accounts for imbalanced sets of false positive and false negative counts (see the methods section for the F1 formula used). This was particularly important given that the methods varied widely in terms of relative fractions of false positive and false negative expression estimates for TEs. F1 accuracy scores across all benchmarked software packages on both gene and TE annotations are displayed in **Figure 3**. In general, TEsingle’s accuracy on gene annotations is comparable to or better than that of popular gene-only single-cell and single-nucleus software packages **(Figure 3A)** with an F1 score of 0.87 for genes on the whole-cell intron retention model, compared to 0.81 for STARsolo-TE, 0.79 for CellRanger-TE, 0.94 for soloTE, and 0.66 for scTE. Thus, including TEs in the gene expression estimation packages does not erode gene estimates, and may improve estimates of gene expression.

**Figure 3:**
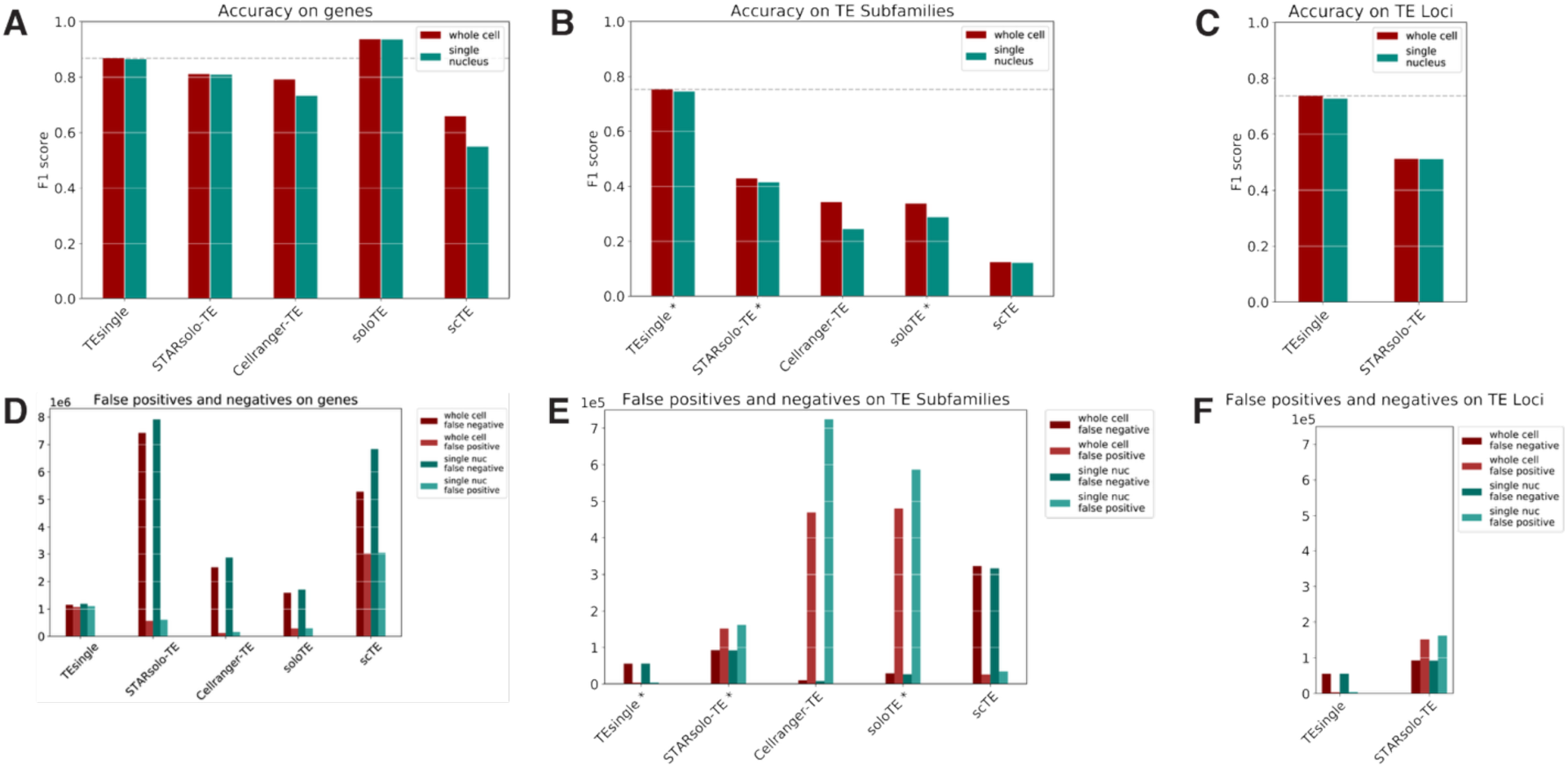
Benchmarking accuracy results. **A-C:** F1 score measures of accuracy for each software benchmarked on simulated whole-cell and snRNA-seq data. **A:** Accuracy measures on gene annotations. **B:** Accuracy measures on TE annotations, by subfamily. Software results denoted with * were summed by subfamily for comparison. **C:** Accuracy measures on TE annotations, by locus, for the two locus-specific software. **D-F:** Plots of the number of annotations measured as false positive (zero expression simulated, nonzero expression measured) and false negative (zero expression measured, nonzero expression simulated). **D:** False positive and negative plots for all benchmarked software on gene annotations. **E:** False positive and negative plots for all benchmarked software on TE annotations, counted by subfamily. **F:** False positive and negative plots for all benchmarked software on TE annotations, counted by locus.

For TEs, we benchmarked locus-specific expression estimates for those TE software packages capable of evaluating TE expression at each the individual locus level (TEsingle and STARsolo-TE). However, most packages combine annotated TE loci into their respective subfamilies and estimate expression at the subfamily level for some or all expressed TEs (Cellranger-TE, soloTE, scTE). Thus, to enable a fair comparison across all packages, we additionally summed the results of the locus-specific TE packages into subfamily-level expression estimates. As such, we display accuracy (F1) measures for TE expression estimates in two separate categories: one that aggregates all expressed TEs by subfamily (**Figure 3B**, TE subfamilies) and one that measures F1 scores at single-locus resolution (**Figure 3C**, TE loci). At the subfamily level (**Figure 3B**), TEsingle showed the highest F1 scores as compared to all TE analysis packages, with F1 scores of 0.75 for TEsingle, 0.51 for STARsolo-TE, 0.34 for CellRanger-TE, 0.34 for soloTE, and 0.12 for scTE. At the locus-specific TE level (**Figure 3C**), TEsingle also outperformed STARsolo-TE with F1 scores of 0.74 for TEsingle and 0.51 for STARsolo-TE. Scores for simulated data generated from the single-nucleus (intron-retention) model are plotted in green alongside these accuracy scores in **Figures 3A-C**, showing very similar relative performance in accuracy at the subfamily level and locus-specific level, respectively. F1 score values for all packages for both single-cell and single-nucleus simulated datasets are available in **Supplemental Table S2**.

Our benchmarking results demonstrate that TEsingle outperforms both gene-focused and TE-focused software packages in accurately evaluating expression of transposable element loci, whether the results are locus-specific or aggregated by subfamily. STARsolo-TE, a slight modification of the STARsolo package (see methods), had the second-best performance on both gene and TE estimation. STARsolo-TE was also the only other software package able to return accurate TE estimates at both the subfamily and locus-specific level. For all packages, the largest contributor to error in gene/TE estimation was false positive and false negative calls, as displayed in **Figure 3D-F**. Many packages showed a tendency to mis-assign gene-derived reads to TEs, resulting in reciprocal false negative calls for genes and false positive calls for TEs. Since there are many more non-expressed TE loci than there are expressed genes and TEs, it was also possible for many packages to have high TE false positive rates without a proportional change in gene false negative rates – for example, by calling the wrong TE locus or subfamily as expressed or non-expressed.

As expected, all TE analysis packages showed poorer performance on single-nucleus synthetic data compared to whole-cell simulations, since single-nucleus simulations contained more intron-spanning gene reads that can be mistaken for expression from intronic TEs. These differences in accuracy between whole-cell and single-nucleus expression estimates can best be seen in the graphs of false positive and false negative rates (**Figure 3D-E**) where single-nucleus false calls (green bars) are typically higher than whole-cell false call rates (red bars). TEsingle achieves high overall accuracy by returning low false positive and false negative rates for both gene and TE expression without compromising its ability to correctly call expressed TEs.

### 4. Applying TEsingle to post-mortem single-nucleus RNA-seq data demonstrates the importance of combining gene and TE analysis

Given TEsingle’s improved accuracy in quantifying gene and TE expression at both the single-cell and single-locus level, we next demonstrated its ability to identify TE expression in single-nucleus sequencing profiles from SNpc tissues of PD patients and age-matched controls. For this showcase, we chose a publicly available postmortem PD snRNA-seq dataset, published by Martirosyan^55^ and colleagues, which had exceptionally good capture of the DA neurons known to degenerate in PD^56^. The specificity of tissue sampling and recovery of many DAs gave sufficient power to do meaningful differential expression (DE) analysis at a cell subpopulation level and reduces background sampling of nearby brain regions. This allowed us to determine whether there was any evidence for elevated TE expression within specific cell types and cell subpopulations in PD.

As demonstrated in **Figure 4A**, the pipeline for snRNA-seq analysis begins with standard alignment and pre-processing steps (see methods), followed by gene and TE expression estimation with TEsingle. TEsingle’s cell-by-gene output table is designed for easy incorporation into standard downstream analysis pipelines, such as Seurat^57^. Using Seurat, we integrated all PD and control samples using Canonical Correlation Analysis (CCA) and identified cell clusters using the Leiden algorithm. These Leiden clusters, visualized by UMAP in **Figure 4B**, revealed seven broad cell types present.

**Figure 4:**
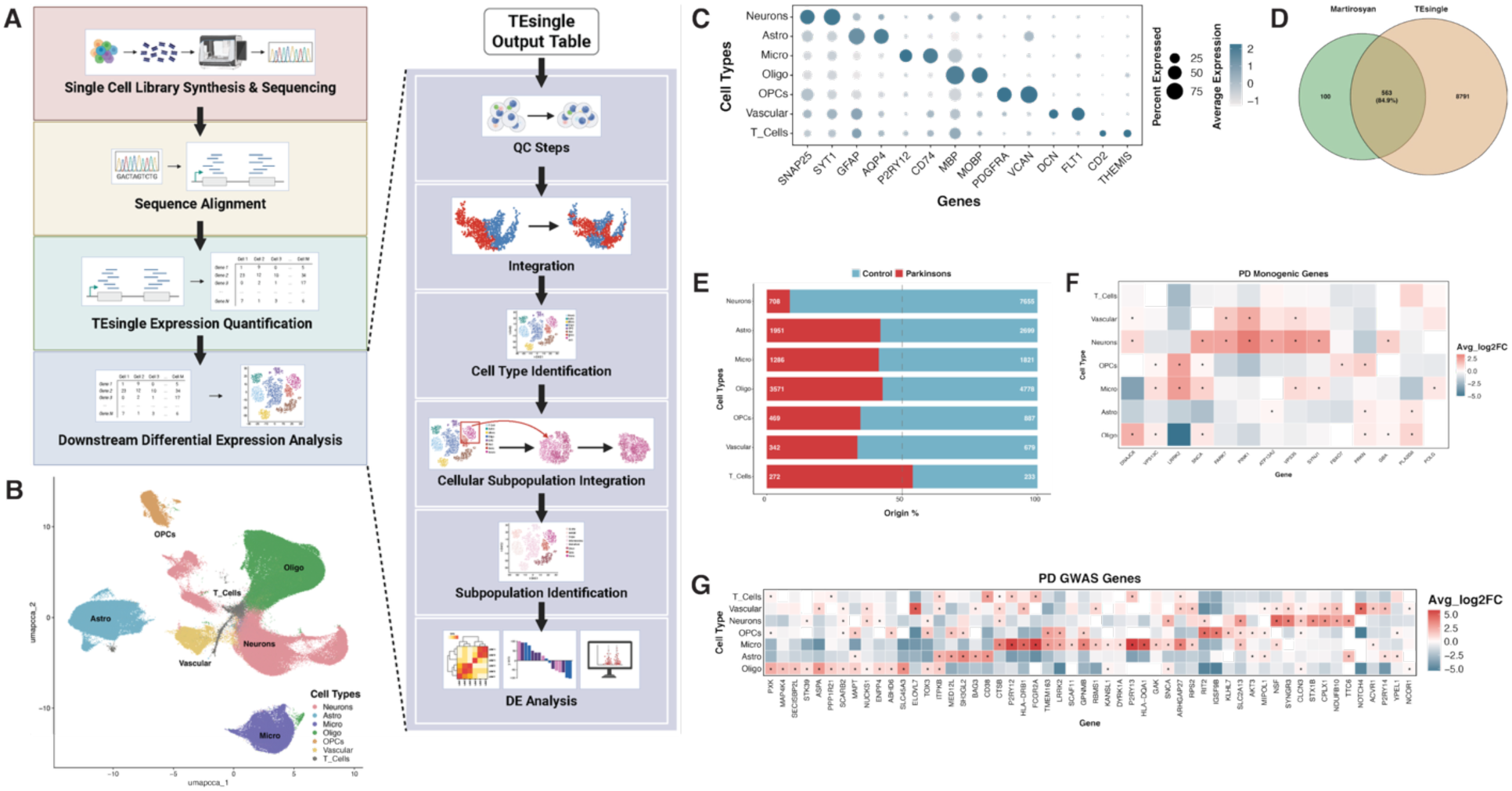
TEsingle enables accurate expression quantification of genes while also providing TE-related information. **A:** A flow chart of Seurat’s downstream analysis workflow of snRNA-seq data to identify differentially expressed genes and TEs after subpopulation identification. **B:** UMAP based Leiden clustering of broad cell types identifying seven categories of cells: Neurons, Astrocytes (Astro), Microglia (Micro), Oligodendrocytes (Oligo), Oligodendrocyte progenitor cells (OPCs), Vascular cells (Vascular), and T-cells. **C:** Dot plot showing the expression of selected marker genes used to identify each broad cell type cluster. **D:** Venn diagram showing the number of DEGs found to be the same between our differential expression analysis and Martirosyan’s analysis. We were able to recapitulate 563 genes (84.9% overlap). **E:** Horizontal bar plot showing the normalized cell counts by sample, where the mean number of cells per sample were counted for each cell type. **F:** Enriched expression of broad cell type populations of selected PD-associated monogenic genes. The average log2FC of normalized counts are shown with significance denoted with * if adjusted p-value (FDR) < 0.05. **G**: Enriched expression of broad cell type populations of selected genes near PD-associated variants based on Martirosyan’s MAGMA analysis on PD GWAS data. The average log2FC of normalized counts are shown with significance denoted with * when adjusted p-value (FDR) < 0.05.

Using the same canonical cell-type-specific marker genes as the Martirosyan group, we were able to confirm the identity of each cluster where broad cellular populations were marked by the following features **(Figure 4C)**: neurons (*SNAP25, SYT1*), astrocytes (Astro) (*GFAP, AQP4*), microglia (Micro) (*P2RY12, CD74*), oligodendrocytes (Oligo) (*MBP, MOBP*), oligodendrocyte progenitor cells (OPCs) (*PDGFRA, VCAN*), vascular cells (*DCN, FLT1*), and T-cells (*CD2, THEMIS*). A complete summary of metadata features for the samples analyzed and the total numbers of recovered nuclei across the broad cell types and subpopulations after quality control can be found in **Supplemental Table 7**.

To ensure that these gene expression-based results from TEsingle were reproducing those previously obtained with a gene-only analysis, we next sought to understand whether we could reproduce the major findings from the original study with TEsingle’s output. DE analysis was performed using the logistic regression method within Seurat to identify differentially expressed genes (DEGs) between PD and control cells within each cell type and/or subpopulation. Of the 663 genes originally identified as differentially expressed in PD cell types, we recapitulated 563 genes (84.9% overlap, **Figure 4D**), despite using different DE statistical methods and incorporating TEs into the analysis. In addition, after normalizing cell counts by sample, where the mean number of cells per sample was calculated for each cell type, we found that most cell types were depleted in PD compared to controls, except for T-cells **(Figure 4E).** We also reproduced the original study’s findings on cell type specific expression of PD risk genes for both monogenic and GWAS associated genes. Using the same high-confidence monogenic genes associated with PD^49,55^, we recapitulated the enrichments for neuronal and glial biased expression in these genes as previously reported in Martirosyan et al.

Specifically, *SNCA* was enriched in neurons and microglia, *DNAJC6* in neurons and oligodendrocytes, and *LRRK2* in microglia and OPCs (**Figure 4F**). With TEsingle’s ability to capture more transcripts for assignment, we were also able to observe some more nuanced gene enrichment in specific cell types, with *SNCA* also significantly upregulated in OPCs and oligodendrocytes, and *DNAJC6* in vascular cells. Several other monogenic PD genes that are enriched within neurons include *PARK7*, *PINK1*, *ATP13A2*, *VPS35*, and *SYNJ1* with added enrichment in vascular cells for *PARK7* and *PINK1*, vascular cells and microglia for *VPS35*, astrocytes for *ATP13A2*, and microglia for *SYNJ1* (**Figure 4F**). We also replicated the original findings for expression of genes located near PD-associated variants identified by their Multi-marker Analysis of GenoMic Annotation (MAGMA) analysis on PD GWAS^55^. TEsingle recapitulates cell-type biased expression of *SNCA* in neurons, *P2RY12* in microglia, and *LRRK2* in microglia and OPCs (**Figure 4G**). Microglia and oligodendrocytes also showed relatively high levels of *SNCA* but were not as specifically elevated for SNCA expression as the neurons (by log2FC across cell types). These results confirmed that we were able to reproduce major findings from a gene-only analysis using TEsingle’s output and show that there are specific cell type biases in expression level of PD-associated monogenic and GWAS genes. A complete list of all DE genes and TEs used to make **Figure 4F and 4G** can be found in **Supplemental Table 8**. As discussed below, we found thousands of TE loci differentially expressed in PD tissues as compared to controls, accounting for the majority of the DE features captured by TEsingle but not identified in the original analysis.

### 5. PD neurons show elevation of young and human-specific TEs

A hallmark of PD development includes neuronal stress and neuroinflammation^58^. While inflammatory triggers within PD remain unknown, one emerging mechanism could be the activation of TEs. TE reactivation has been linked to aging and several other neurodegenerative diseases^3,8–10,59–61^, but its role in PD remains understudied^62–67^. This prompted an exploration of TE expression across the cell types and subpopulations present in SNpc tissues. Given that PD is characterized by the progressive loss of DA neurons, we first asked whether TE expression would be detectable and differentially abundant across DA neurons from PD patient samples. Neuronal subpopulations were identified through Leiden re-clustering of all neurons captured, which returned four major neuronal types, as displayed in the UMAP **(Figure 5A).** Using canonical neuronal subtype-specific marker genes and comparing to the cluster marker genes **(Supplemental Table 9A),** we were able to assign neuronal subtypes based upon the following features: DAs (DopaN) (*TH)*, inhibitory neurons (*GAD1)*, and excitatory neurons (*SLC17A6)* **(Figure 5B).** Neurons that were not clearly expressing known markers were labeled as “Other_Neurons”. DE analysis of PD vs control cells within each neuronal subpopulation revealed a general upregulation of TEs within PD-derived DAs, inhibitory, and excitatory neurons **(Figure 5C)**. A complete list of DE genes and TEs between PD and control neuronal subpopulations can be found in **Supplemental Table 10**.

**Figure 5:**
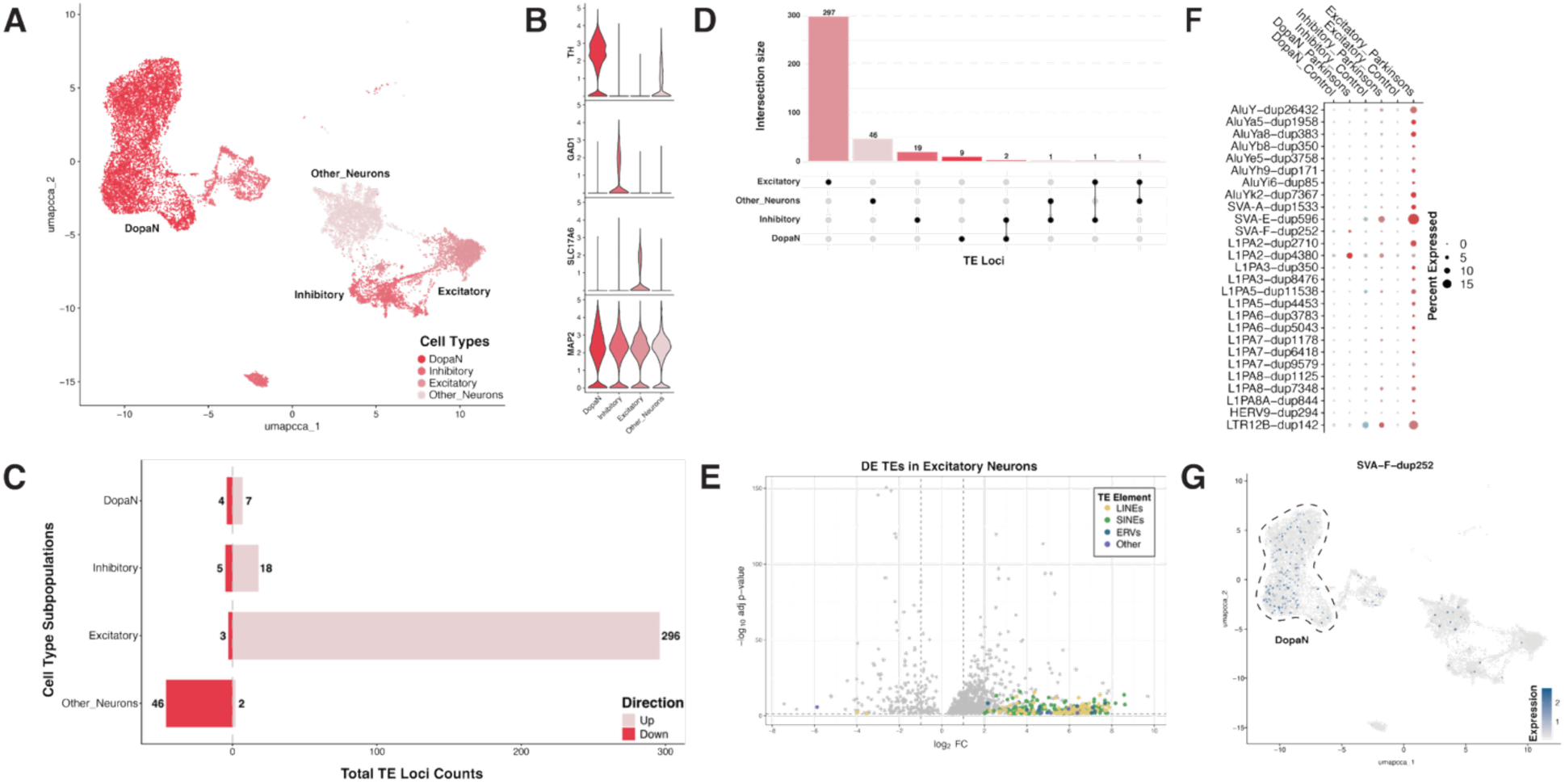
PD patient neurons show elevation of young and human-specific TEs. **A:** UMAP based Leiden clustering of all neurons in the Martirosyan dataset showing the identification of four neuronal subpopulations: dopaminergic (DopaN), inhibitory, excitatory, and other neurons. **B:** Violin plots displaying the marker genes that are enriched within each neuronal subpopulation. **C:** Horizontal bar plot counting the total number of DE TE loci within each PD-derived neuronal subpopulation after differential expression analysis to the matched control neuronal subpopulation. **D:** Upset plot showing the total counts of DE TE loci within each PD-derived neuronal subpopulation or shared between neuronal subpopulations. Counts includes both up and downregulated DE TE loci. **E:** Volcano plot of fold change versus negative log10 adjusted p-values displaying all significantly DE TE loci (colored dots) against all significantly DE genes (gray dots) within PD-derived excitatory neurons. Each colored point is a specific TE locus that is colored based on their TE class, as noted in the legend. **F:** Dot plot showing the percent expression of selected younger and human-specific TE loci between PD and control neuronal subpopulations. The size of the dot represents the percentage of cells that express the TE locus, and the color encodes the diagnosis of PD (red) or control tissue (blue). **G:** Feature plot showcasing the SVA-F-dub252 TE locus that is specifically enriched within dopaminergic neurons. Colored dots represent single cells on the neuron UMAP presented in Figure 5A, where the darker the colored cells, the higher the expression value of the SVA-F-dub252 TE locus within that cell.

When comparing across neuronal subtypes, we found that most of the PD-elevated TEs were unique to a neuronal subpopulation, with only a handful of shared PD-elevated TEs found in more than one neuronal population (see upset plot in **Figure 5D**). We found the largest set of elevated TE loci in excitatory neurons **(Figure 5C, E)**, which showed elevated expression of hundreds of LINE, SINE, and ERV type TEs. Many of these elevated TEs belonged to relatively young, intact and/or human-specific TE subfamilies (**Figure 5F**), such as a young LINE element (L1PA2-dup4380), sequences from endogenous retrovirus families (HERV9-dup294,LTR12B-dup142), and from human-specific SVA elements (SVA-E-dup596, SVA-F-dup252). In particular, several of these young and intact PD-elevated TEs were specific to DA neurons, such as SVA-F-dup252, which is both PD-elevated and expressed nearly exclusively in DA neurons **(Figure 5G)**. SVA elements are a composite of other TEs (SINE-VNTR-Alu elements) that are still transcriptionally active in humans and can use LINE-1 machinery for retrotransposition^68,69^. Other related SVA insertions have been previously associated with human diseases^70–76^, and in particular, a related SVA-F locus has been associated with X-linked dystonia-parkinsonism^77^. Altogether, this demonstrates that relatively young and intact TEs are elevated in specific neuronal subpopulations of PD patient tissues, and that some of the more interesting TE candidates are specific to DA midbrain neurons, the cell type most vulnerable in PD.

### 6. PD tissues are enriched for a population of Reactive Astrocytes marked by young human-specific TEs

Next, we investigated TE elevation within astrocytes to assess if TE expression can be correlated to a specific subpopulation/state. Astrocytes have numerous roles in maintaining neuronal health and play critical roles in mediating neuroinflammatory processes within the central nervous system^78^. In PD, the presence of pro-inflammatory astrocytes has been associated with signs of neuroinflammation^79^, and we wondered if pro-inflammatory astrocyte states would also be associated with an enrichment of TE expression. Astrocyte subpopulations/substates were identified through Leiden re-clustering and cluster marker expression, with six astrocyte subpopulations present in the SN tissues **(UMAP**, **Figure 6A).** By looking at cluster marker gene expression **(Supplemental Table 9B),** and comparing with previously published datasets of astrocyte states^55,80–82^, we were able to identify these astrocyte subpopulations as: homeostatic astrocytes (HAs), intermediate astrocytes (InAs), a subpopulation of astrocytes that are in between homeostatic and reactive astrocytes characterized by Serrano-Pozo et al^81^, reactive astrocytes (RAs) (*HSPH1)*, disease-associated astrocytes (DAAs) (*ADAMTSL3)*, inflammatory astrocytes (IfAs) (*C3*), and metallothionein expressing astrocytes (MAs) (*MT1G)* **(Figure 6B)**. TEsingle expression analysis did not reveal a global elevation of TE loci within PD-derived astrocytes.

**Figure 6:**
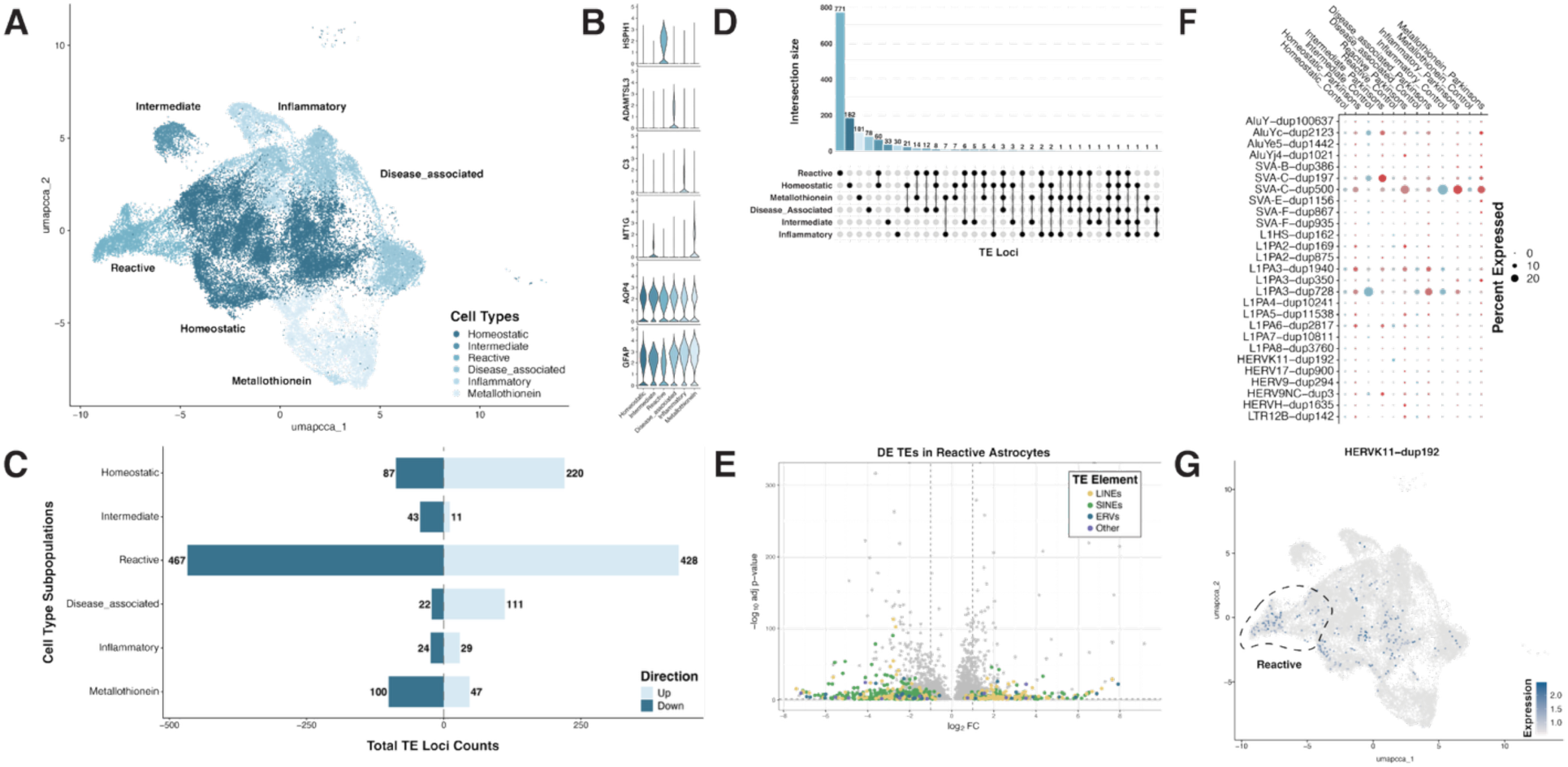
PD tissues are enriched for a population of reactive astrocytes marked by young human-specific TEs. **A:** UMAP based Leiden clustering of all astrocytes in the Martirosyan dataset showing the identification of six astrocytic states: homeostatic, intermediate, reactive, disease associated, inflammatory, and metallothionein. **B:** Violin plots displaying the marker genes that are enriched within each astrocytic state. **C:** Horizontal bar plot counting the total number of DE TE loci within each PD-derived astrocytic state after differential expression analysis to the matched control astrocytic state. **D:** Upset plot showing the total counts of DE TE loci within each PD-derived astrocytic state or shared between astrocytic states. Counts includes both up and downregulated DE TE loci. **E:** Volcano plot of fold change versus negative log10 adjusted p-values displaying all significantly DE TE loci (colored dots) against all significantly DE genes (gray dots) within PD-derived reactive astrocytes. Each colored point is a specific TE locus that is colored based on the TE class they belong to, as noted in the legend. **F:** Dot plot showing the percent expression of selected younger and human-specific TE loci between PD and control astrocytic states. The size of the dot represents the percentage of cells that express the TE locus, and the color encodes the diagnosis of PD (red) or control tissue (blue). **G:** Feature plot showcasing the HERVK11-dup192 TE locus that is specifically enriched within reactive astrocytes. Colored dots represent single cells on the astrocyte UMAP presented in Figure 6A, where the darker the colored cells, the higher the expression value of the HERVK11-dup192 TE locus within that cell.

Instead, we found that PD-derived reactive astrocytes (RAs) had the largest number of dysregulated TE loci **(Figure 6C)** with the vast majority of the DE TEs being unique to that astrocytic state **(Figure 6D)**. Of the PD dysregulated TEs within the RA subpopulation, we found representatives of each of the major LINE, SINE, and LTR classes of TEs **(Figure 6E)**. A complete list of DE genes and TEs between PD and control astrocyte subpopulations can be found in **Supplemental Table 11**.

When comparing across astrocytic states, we found a number of relatively young TEs that were commonly elevated in PD **(Figure 6F)**. For example, SVA-C-dup500 was elevated in PD RAs as well as IfAs, and MAs. In contrast, we also looked for TEs that were unique to specific astrocytic subpopulations, where we found several young human endogenous retroviruses, such as HERVK11-dup192, that uniquely marked the RA cluster **(Figure 6G)**. HERVK11 is a part of the endogenous retrovirus (ERV) superfamily that makes up of 8% of the human genome^83^. Additionally, HERVK11 is part of the evolutionarily youngest human-specific ERV family^84^. Dysregulation of HERVK supergroup expression has been implicated in several neurodegenerative diseases, including ALS and Alzheimer’s Disease^8,85–89^, and has been shown to be activated by inflammatory responses mediated by interferon regulatory factor 1 (IRF1) and NF-kappa-B signaling in astrocytes and neurons in ALS^90^. Although not many studies have focused on the role of HERVKs in PD, one whole genome sequencing analysis has found evidence for a polymorphic HERVK element that may contribute to PD development^91^. Together, the inflammatory responsiveness of HERVK elements and the broader involvement of polymorphic HERVK in PD suggest a potential role for HERVK11-dup192 within RAs in PD-related neuroinflammatory processes.

### 7. Chemokine Response Microglia are enriched in PD and marked by young and human-specific TE loci

Given the well-established roles for microglia in driving neuroinflammatory cascades^92^, we next sought to investigate whether specific microglial subpopulations are correlated with increased TE activation in PD. Microglial states were identified through Leiden re-clustering of all microglia, with five microglial states identified in the PD and control SNpc tissues **(Figure 7A).** Cluster marker genes **(Supplemental Table 9C),** and previously published markers of microglial states^55,93,94^ helped us characterize each of these microglial clusters **(Figure 7B)**: homeostatic microglia (HMs), cytokine responding microglia (CRMs) (*DUSP1*), interferon responding microglia (IRMs) (*DDX21*), antigen presenting microglia (APMs) (*AIF1*), and disease associated microglia (DAMs) (*TREM2*). DE analysis revealed a global elevation of TEs within microglia **(Figure 7C)** with cytokine response microglia (CRMs) showing the largest set of PD-upregulated TEs **(Figure 7E).** A complete list of DE genes and TEs between PD and control microglial subpopulations can be found in **Supplemental Table 12**.

**Figure 7:**
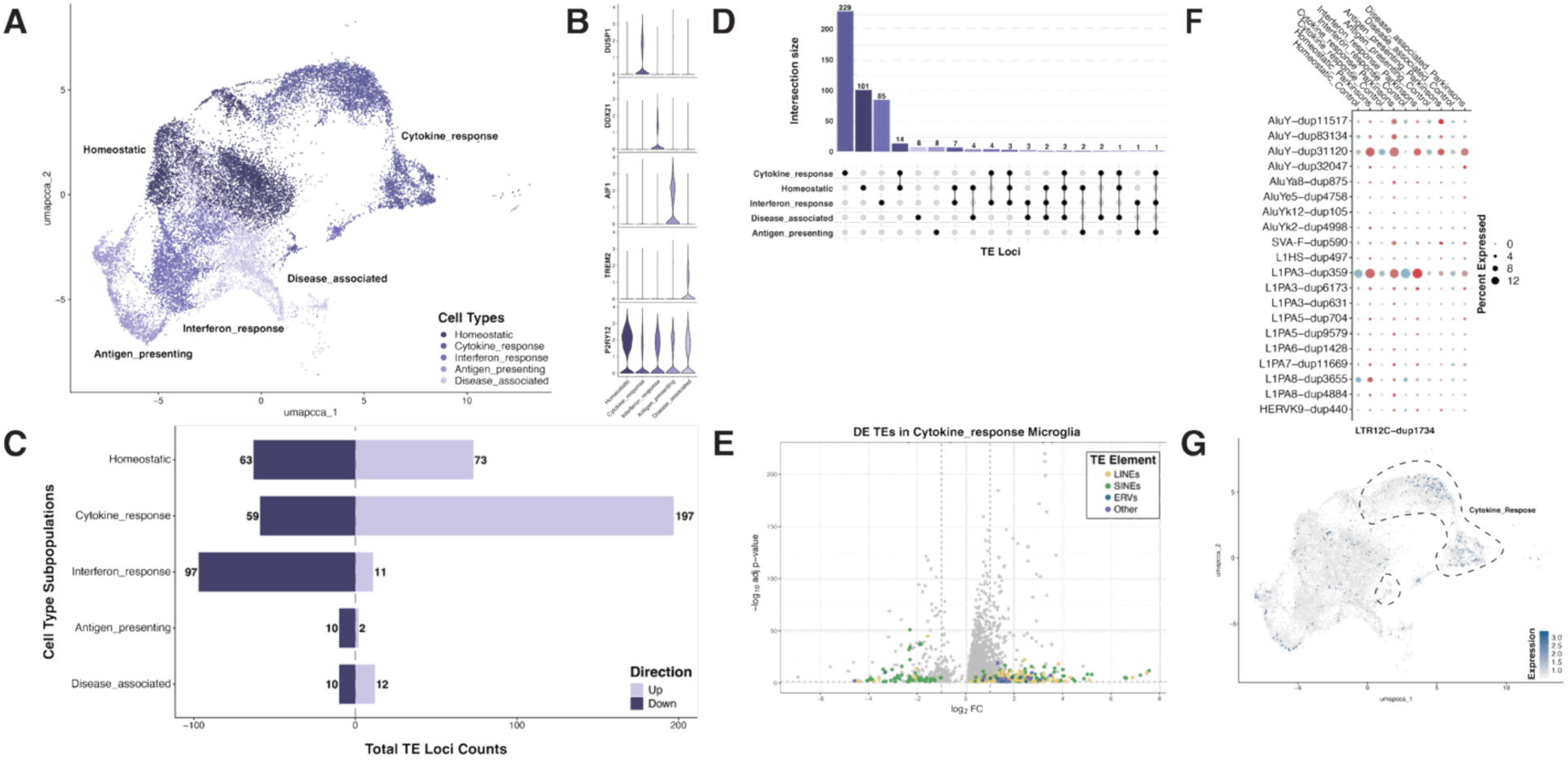
Chemokine response microglia are enriched in PD and marked by young and human-specific TE loci. **A:** UMAP based Leiden clustering of all microglia in the Martirosyan dataset showing the identification of five microglial states: homeostatic, cytokine responding (cytokine_response), interferon responding (interferon_response), antigen presenting (antigen_presenting), and disease associated (disease_associated) microglia. **B:** Violin plots displaying the marker genes that are enriched within each microglial state. **C:** Horizontal bar plot counting the total number of DE TE loci within each PD-derived microglial state after differential expression analysis to the matched control microglial state. **D:** Upset plot showing the total counts of DE TE loci within each PD-derived microglial state or shared between microglial states. Counts includes both up and downregulated DE TE loci. **E:** Volcano plot of fold change versus negative log10 adjusted p-values displaying all significantly DE TE loci (colored dots) against all significantly DE genes (gray dots) within PD-derived cytokine responding microglia. Each colored point is a specific TE locus that is colored based on the TE class they belong to, as noted in the legend. **F:** Dot plot showing the percent expression of selected younger and human-specific TE loci between PD and control microglial states. The size of the dot represents the percentage of cells that express the TE locus, and the color encodes the diagnosis of PD (red) or control tissue (blue). **G:** Feature plot showcasing the LTR12C-dup1734 TE locus that is specifically enriched within cytokine responding microglia. Colored dots represent single cells on the microglia UMAP presented in Figure 7A, where the darker the colored cells, the higher the expression value of the LTR12C-dup1734 TE locus within that cell.

As was seen for neurons and astrocytes, most of the dysregulated TEs were specific to a single microglial state, but we did find a few PD-elevated TEs that were shared between PD HMs and CRMs **(Figure 7D).** Looking specifically at the CRM cluster, we noted that the PD-associated and up-regulated TEs were more likely to derive from LINE and ERV classes of TEs, while SINEs formed the majority of the down-regulated TEs (**Figure E**). The PD-specific elevated TEs included relatively young and intact TE loci **(Figure 7F)**, including members of the human-specific LINE-1 subfamily (L1HS-dup497) as well as members of the human specific HERVK9 subfamily (HERVK9-dup440). Example PD-elevated TEs that are unique to a particular microglial substate include an element from the HERV-9 subfamily (LTR12C-dup1734), which specifically marks CRMs **(Figure 7G)**. LTR12C sequences both drive ERV expression and can also act as cis-regulatory elements that modify host gene expression, with studies demonstrating that around 20% of genomic LTR12C elements carrying active enhancer and/or promoter marks, H3K4me3 and H3K27ac^95^. One particularly interesting study of LTR12C mediated gene regulation showed that activation of LTR12C elements led to the activation of novel gene transcription start sites and LTR-derived immunogenic antigens presented on MHC class I^96^. This suggests the potential for some of these TE elements, such as LTR12C, to have broader impact on cellular function outside of driving TE expression.

Finally, while neurons, astrocytes, and microglia showed the greatest evidence for widespread dysregulation of TEs in PD tissues, we also noted dysregulation of TEs in other cell types including oligodendrocytes (ODC). Information about PD-associated TE expression alterations in ODCs can be found in the Supplemental sections **(Supplemental Figure 2, Supplemental Table 9D, and Supplemental Table 13)**.

## Discussion

TEsingle is a novel gene and TE single-cell software package that uses strategies for expression analysis derived from a deep understanding of how genes and TEs coexist within genomes and how those transcripts are captured and amplified in single-cell assays. Locus-specific resolution of TE expression enables an association of TEs with local chromatin environments, expression relative to adjacent genes, and disambiguation of expressed intronic TEs from overlapping genes. Furthermore, each specific TE locus differs in the number of mutations (single-nucleotide and larger) that constrain its potential impact on cellular function. For example, each TE locus differs in its relative ability to generate RNA transcripts from its own promoter. Furthermore, only a small fraction of TEs can code for intact proteins, generate reverse-transcribed cDNA, or transpose^19,97^. Accurate tracking of TE expression at the locus-specific level enables better prediction of the potential for those transcriptionally active TEs to impact cellular function.

As demonstrated by our benchmarking results in **Figure 3**, most TE expression analysis software suffers from high rates of false positive and false negative calls for both genes and TEs. This was especially true for simulated datasets that contained larger fractions of unprocessed pre-mRNAs, which are common in snRNA-seq experiments.

Furthermore, whole-cell single-cell datasets are difficult to obtain for archival and post-mortem tissue profiling studies, necessitating software that can handle higher levels of intronic reads from pre-mRNAs or intron retention events (e.g., snRNA-seq data). This need for accurately disambiguating gene/TE expression is important for both TE researchers, and for those who want to ensure their gene expression estimates are not confounded by overlapping and interwoven TEs. Thus, accurately handling TEs in any gene expression analysis will improve estimates of gene expression.

Improved methods for TE expression analysis have led to an increased appreciation for the impact of TE activity in aging^2^ and specifically in aging-associated neurodegenerative disorders^98–101^. However, it remains unclear whether elevated TE expression is a simple consequence of aging cells, or whether specific TEs might be particularly associated with individual cell types and cellular states. To demonstrate the ability of TEsingle to track TE expression at a locus-specific and single-nucleus level in real data, we applied TEsingle to postmortem tissue samples from the substantia nigra of Parkinson’s Disease patients. This enabled us to determine that several young TEs from the SVA family are specifically expressed in the dopaminergic neurons of PD patients. Furthermore, we found a global elevation of hundreds of TE loci in the surrounding excitatory neurons, many of which belong to young TE subfamilies at loci that are full-length and/or have coding potential (L1PA2, L1PA3, and HERV-9/LTR12). Looking at surrounding glial cells of the subtantia nigra, we found differences in TE expression that were specific to individual glial substates, with global elevation of TEs among microglia expressing markers associated with a “cytokine response”. In general, TEs were highly cell-type- and cellular-state-specific in their expression patterns, with very little shared TE expression across cell types and states. Finally, cellular states with the largest number of elevated TEs in PD tissues were typically pro-inflammatory glial states, such as “reactive” astrocytes and “cytokine response” microglia. This suggests that specific TE loci may act as inflammatory cell-state specific markers and may add to our understanding of the cellular states adopted by glia in response to pro-inflammatory stimuli. Future work to understand the relative impact of dysregulated TEs in PD may establish whether TEs primarily impact the expression of neighboring genes^63,66^ or whether the expressed TEs themselves might contribute to cell state alterations^62^.

## Methods

### Generating simulated data with FluxSimulator

The GTF annotation file for the gene annotations found in the T2T human genome was obtained from RefSeq^102^ and the GTF file for transposable element annotations was generated from the RepeatMasker^41^ table obtained from the UCSC genome database^103,104^. The TE annotation table was then filtered to remove regions of low complexity, simple repeats, rRNA, scRNA, snRNA, srpRNA, and tRNA. Annotation files prepared in this fashion have been provided for several genomes as indicated at the TEsingle GitHub repository, but TEsingle can also use custom annotations that conform to the specified GTF format.

In order to simulate a scRNA-seq experiment, we first decided to model one with four different groups of cells, each with their own expression profile. We modeled each group as 1250 cells, with each cell having an average of 300,000 transcripts total expressed, from which 10,000 randomly sampled transcripts were used to generate cDNA. Each transcription product was assigned a cell barcode and randomly generated UMI, which was then used to generate sequencing files for the matching barcode and UMI for each read simulated from the library later. From 10,000 expressed transcripts for each cell, we then simulated all steps of cDNA synthesis, amplification, and sequencing to generate 100,000 reads for each cell using FluxSimulator^105^ (version 1.2.3). These simulated libraries were built both with and without retained introns, so that the percentage of processed and unprocessed transcripts could be fine-tuned in the final sequence files. All simulated data reads were mapped back to the T2T genome using STAR^50^ (version 2.7.11b) and any reads that could not be aligned to the annotation of origin within a maximum of 100 multi-mapping locations were discarded.

Reads were then aligned to the genome using STAR to generate a SAM file of all aligned reads. Transcripts were then chosen from either the fully spliced or fully unspliced transcript files to match the intron retention fractions for whole-cell or snRNA-seq data, and all reads generated from those transcripts could be combined into one fastq file. Using the known transcript of origin information, we could then generate a ground truth file of expressed transcript counts that could be observed from each fastq dataset.

### Estimating intron retention in single-cell and single-nucleus expression data

The likelihood of a cDNA being from an intronic region of an unspliced transcript, given that we know the transcript is unspliced, is proportionate to the fraction of bases of that transcript that are intronic. But the likelihood of any cDNA being from an unspliced transcript to begin with is the fraction of transcripts that are unspliced in the sequenced pool. Thus, the fraction of unspliced transcripts represented in a single-cell or single-nucleus RNA-seq experiment is equal to the fraction of total coverage of purely intronic sequence divided by the fraction of sequenced bases that are intronic:

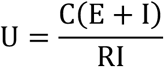

where U is the fraction of transcripts in the sequencing run that were unspliced, C is the number of total bases of intronic coverage in the sequencing run, E is the number of bases of exonic sequence evaluated, I is the number of bases of intronic sequence evaluated, and R is the total number of bases of read coverage sequenced over both intronic and exonic regions.

First a cleaned .bed file of regions that are known to be within the gene transcripts, but never part of exons or TEs, is generated using BEDtools^106^. Then a cleaned .bed file of regions known to be within the gene exons, but never part of introns or TEs, is generated. Total bases encompassed by each of these files is calculated. Using BEDtools, we can then calculate the coverage over each base represented by those .bed files and then sum those coverages to get the total coverage over the introns, exons, or both.

Intron retention fractions were thus estimated from single-cell^48^ and single-nucleus^49^ RNA-seq experiments.

### Single-cell/nuclei RNA-seq data pre-processing and alignment, and parameters for running related software

To prepare sequencing files to be run by TEsingle, the sequence files must first be aligned. We recommend using STARsolo with the following settings:

--alignIntronMax 1000000

--alignIntronMin 20

--alignMatesGapMax 1000000

--alignSJDBoverhangMin 1

--alignSJoverhangMin 8

--outFilterMismatchNmax 999

--outFilterMismatchNoverReadLmax 0.04

--outFilterMultimapNmax 100

--winAnchorMultimapNmax 200

--outFilterType BySJout

--outSAMattributes NH HI AS nM CR CY UR UY CB GX GN sS sQ sM

--outSAMheaderHD @HD VN:1.4

--outSAMstrandField intronMotif

--outSAMtype BAM SortedByCoordinate

--sjdbScore 1

--soloUMIlen 12

--soloType CB_samTagOut

--soloCellFilter EmptyDrops_CR 5000 0.99 10 45000 90000 500 0.01 20000 0.01 10000

--soloCBmatchWLtype 1MM

--soloUMIdedup 1MM_All

We have compared F1 scores for allowing each read to align only once, up to ten times, one hundred times, and one thousand times (see **Supplemental Figure S1A** for a plot comparing accuracy in calling gene annotations, and **Supplemental Figure S1B** for a plot comparing accuracy in calling TE annotations, or **Supplemental Table S3** for full F1 scores). From this comparison we have concluded that allowing reads that map up to 100 times provides the optimal balance of speed and accuracy on our datasets.

### Benchmarking software comparison settings

#### STARsolo-TE

To fairly compare STARsolo with TEsingle on simulated data with intron retention, we ran STARsolo with the following settings:

--outFilterType BySJout

--outSAMstrandField intronMotif

--alignSJoverhangMin 8

--alignSJDBoverhangMin 1

--alignIntronMin 20

--alignIntronMax 1000000

--alignMatesGapMax 1000000

--sjdbScore 1

--soloType CB_UMI_Simple

--soloCBmatchWLtype 1MM_multi_Nbase_pseudocounts

--outSAMattributes NH HI AS nM CR CY UR UY CB UB GX GN sS sQ sM

--soloFeatures GeneFull

--clipAdapterType CellRanger4

--outFilterScoreMin 30

--soloCellFilter EmptyDrops_CR 3000 0.99 10 45000 90000 500 0.01 20000 0.01 10000

--outFilterMultimapNmax 100

--winAnchorMultimapNmax 150

--outFilterMismatchNmax 999

--outFilterMismatchNoverReadLmax 0.04

--soloMultiMappers EM

--outSAMtype BAM SortedByCoordinate

--outSAMheaderHD @HD VN:1.4

--soloUMIfiltering MultiGeneUMI_CR

--soloUMIdedup 1MM_CR

--soloUMIlen 12

The STAR index built for running STARsolo on transposable elements required a modified annotation file, in which every TE locus was given both a unique transcript_id and a unique gene_id and then appended to the appropriate file of gene annotations. If a unique gene_id is not assigned to each TE locus then index-building will fail, as otherwise, a TE subfamily could be construed as separate transcript exons of the same gene, which would then span multiple chromosomes. This change means that STARsolo results are locus-specific, and so subfamily-level count tables were generated by adding counts assigned to each locus of a single subfamily for each cell, to aggregate by subfamily.

#### CellRanger-TE

To compare TEsingle accuracy with that of CellRanger, we used the CellRanger-TE method outlined in^59^ to generate count tables for the simulated whole-cell and single-nucleus RNAseq data. For that method, to estimate TE expression, a single initial annotation GTF file must be made containing the annotations for all genes and all TE loci. If each TE locus of a given subfamily shares gene_id then any reads that align to multiple loci within the subfamily are given the same alignment annotation, meaning the CellRanger pipeline can then return subfamily-level counts of TE expression. This modification of the original gene annotation file can potentially lead to reads aligning to intronic TEs being discarded if they align with more than one subfamily, or being biased towards being assigned to the TE subfamily instead of the surrounding gene if the read is fully within the intron of the gene, where if they had been compared only to the gene annotations they would have been fully assigned to the gene even if they originated from expression of the intronic TE and not the surrounding gene. To fairly compare CellRanger-TE with TEsingle we set used the following settings at the command line when running CellRanger-TE:

--include-introns=true

--chemistry=auto

#### scTE

The scTE software pipeline^52^ was published in 2021 to assess for TE expression in single-cell RNA sequencing data. This pipeline resolves reads that map to multiple TE loci by restricting the alignment step to report only the highest scoring alignment for each read, regardless of how many locations within the genome it aligns to. The scTE software then takes in a file of reads aligned to the genome and files for both gene and TE annotations, then by default assigns reads first to gene exons and then to TE subfamilies that overlap with the highest scoring alignment.

To estimate the accuracy of the count tables produced by the scTE pipeline we aligned the simulated whole-cell and snRNA-seq data using the settings recommended for using STARsolo to generate an alignment file for scTE:

--winAnchorMultimapNmax 100

--outFilterMultimapNmax 100

--outSAMmultNmax 1

We then used the default scTE parameters for 10x to generate the expression count matrix.

#### soloTE

soloTE is a software pipeline^53^ published in 2022 to assess for TE expression in single-cell RNA sequencing data, with the capability of reporting counts for TEs at the locus level for reads that meet a minimum MAPQ threshold (by default soloTE reports counts at the locus level for reads that only map uniquely to the genome), and to report counts for TEs at the subfamily level for any reads that fail to meet that threshold.

To estimate the accuracy of the count tables produced by the soloTE pipeline we first aligned the simulated whole-cell and snRNA-seq data using the settings recommended for using STARsolo to generate an alignment file for soloTE:

--winAnchorMultimapNmax 100

--outFilterMultimapNmax 100

--outSAMmultNmax 1

Count tables produced by soloTE were aggregated at the subfamily level, for all TEs, before accuracy metrics were calculated because soloTE reports TE counts as a mixture of subfamily-level counts and locus-specific counts, depending on the MAPQ score for each read. We felt that aggregating locus-specific counts with their corresponding subfamily counts would make for the most comparable results for soloTE to all software tested.

#### TEsingle

TEsingle is run on the command line and requires Python (version 3.2 or greater) and the following packages: pysam (version 0.9 or greater), networkx, scipy, and numpy. For input, it takes an aligned BAM or SAM file, one annotation file for genes in GTF or GFF format, and one annotation file for TEs in GTF or GFF format. We recommend aligning sequence data using STARsolo with the settings listed in this paper, but for other aligners the barcode should be stored in the ‘CB’ tag, and the UMI in the ‘UR’ tag in the BAM file. TEsingle expects that correction of cell barcodes to the whitelist has already taken place during the alignment step but also allows for a threshold for the minimum number of UMIs for a cell barcode to be counted (by default this is set to 1000 and can be changed using the --cutoff argument). Cleaned annotation files for genes and TEs for several model organisms and assemblies have been previously generated and shared in the supporting documentation for our previous software, TEtranscript.

TEsingle can take in a novel gene and/or TE annotation file, so long as the annotation files are consistent with the genome sequencing files used for alignment, and contain the requisite genomic location, element name, and gene_id for every gene transcript, plus transcript_id, family_id, and class_id for every TE transcript.

### F1 score calculation

For each genetic element, the count estimated by the software evaluated was classed as ‘assessed accurately’ if the count measured for expression within a given cell was within 15% of the underlying simulated number of transcripts for that genetic element. A count was classed as ‘false positive’ if the count estimated was >0 while the underlying truth was that zero transcripts had been simulated from that genetic element. Similarly, a count was classed as ‘false negative’ if the count estimated was 0 while the underlying truth was that one or more transcripts had been simulated from that genetic element. Counts were classed as ‘overcount’ if the genetic element was simulated to have expression and was measured to have an expression >115% of the ground truth expression simulated and classed as ‘undercount’ if they were measured to have an expression <85% of the ground truth expression simulated. For each simulated sequencing experiment then the total number of counts classed as false negative, overcount, undercount, and accurate will equal the total number of genetic elements simulated to have expression for each cell within the experiment, but there is no limit on the number of counts that could be classed as false positives.

For these classifications we can then define a software’s precision as the measure of how many genetic elements were assessed accurately out of all elements measured to be expressed:

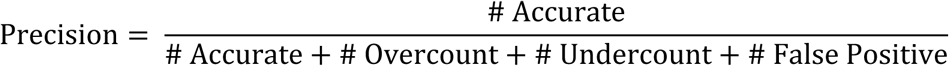

We can further define a software’s sensitivity as the measure of how many genetic elements were assessed accurately of all elements simulated to have expression:

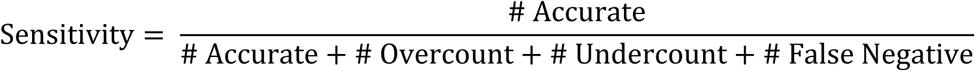

The F1 score is then the harmonic mean of both precision and sensitivity:

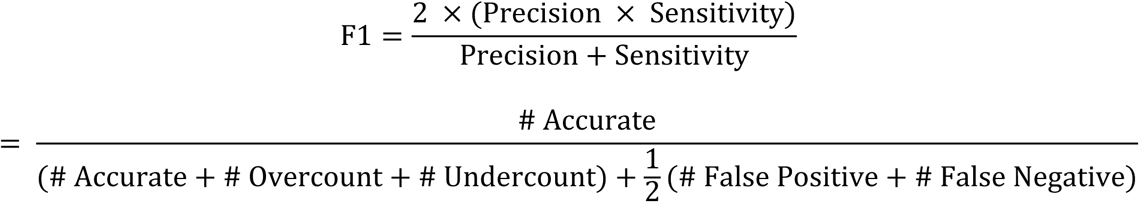

With these definitions, we can then classify the count results for each annotation expressed or measured in each simulated cell and calculate an F1 score for each software on each simulated dataset.

### Preprocessing & Quality Control of Martirosyan snRNA-seq Data for TEsingle

Fastq files generated by Martirosyan and colleagues^55^ were aligned with STAR^50^ against the T2T (T2T-CHM13v2.0) genome^107^ using the parameters described above. TEsingle is then run against the T2T gene (JHU RefSeqv110 + Liftoff v5.2) and TE annotations (RepeatMasker v4.1.2p1.2022Apr14) obtained from the CHM13 GitHub repository^108^, filtering out cell barcodes associated with less than 1000 UMI. After quantification with TEsingle, “empty” cells were filtered using EmptyDrops^109^ with an FDR threshold of 0.1%. DoubletFinder^110^ was run to identify and remove doublets in the dataset, using the multiplex rate provided by 10x Genomics^111^. The cells that passed the QC filters from all samples were then combined into a single MEX file with corresponding feature and barcode sequences and loaded into Seurat^57^ (version 5.3.0) with associated metadata^55^.

### Integration and Clustering of snRNA-seq Data

The Martirosyan dataset was split by diagnosis (PD or control) and counts were log-normalized and linearly transformed. Principal component analysis (PCA) was performed using the top 3000 highly variable features, calculating up to 30 PCs. The dataset was then integrated using Anchor-based CCA integration with gene-only features. Cluster identification was performed using KNN-graph and Leiden algorithm (resolution of 0.5). Dimensional reduction using UMAP^112^ was performed on the integrated data for visualization.

### Cell Cluster Identification and Annotation of snRNA-seq Data

To identify broad cell types, we used the following list of cell type-specific markers previously described by Martirosyan and colleagues^55^: Neurons (*SNAP25, SYT1*), astrocytes (*GFAP, AQP4*), microglia (*P2RY12, CD74*), oligodendrocytes (*MBP, MOG*), oligodendrocyte precursor cells (OPC) (*PDFGFRA, VCAN*), pericytes/endothelial/VLMC (*DCN, FLT1*) and T-cells (*CD2,THEMIS*).

### Differential Analysis of PD-associated Genes in Broad Cell Types

Differential analysis was performed to identify differentially expressed genes and TE loci from one broad cell population between the other broad cell populations using a logistic regression framework. Genes and TE loci were considered differentially expressed if adjusted p-value (FDR) < 0.05.

### Analysis of Neuronal and Glial Cell Subpopulation

Neurons, astrocytes, microglia, and oligodendrocytes were extracted from the integrated Martirosyan dataset, split by diagnosis, and re-normalized using SCTransform^113^ with default settings. Integration, cluster identification and UMAP dimensional reduction were performed as previously described. Cluster markers were identified using a logistic regression framework to determine differentially expressed genes (log fold change threshold of 0.5 and minimum percentage of expressed cells of 0.1). We also used the following cell type-specific markers to identify specific neuronal subpopulations (*TH* for DAs, *GAD1* for GABAergic/inhibitory neurons, and *SLC17A6* for glutaminergic/excitatory neurons). Astrocyte, microglia, and oligodendrocyte subpopulations were identified using the following markers: astrocytes (*HSPH1, ADAMTSL3, C3*, *MT1G, AQP4*), microglia (*DUSP1, DDX21, AIF1, TREM2*), and oligodendrocytes (*OPALIN, OLIG1, DNAJC3, HMOX2*). Cluster annotation was replicated largely following how Martirosyan and colleagues named each subpopulation. Subpopulations were then further identified using additional previously published datasets^55,81,93,114^.

### Differential Analysis of TE Loci and Genes in Cell Subpopulations

Differential analysis was performed to identify differentially expressed genes and TE loci from the same subpopulation of cells between PD and control tissues, using a logistic regression framework. Genes and TE loci were considered differentially expressed if adjusted p-value (FDR) < 0.05.

## Supplemental Table Legends

Supplemental Table 1: Inferred intron retention rates across single-cell and single-nucleus RNA-seq data.

Intron retention fraction calculated from sequencing data for each sample specified. Whole-cell intron retention fraction was approximated as 0.2 for simulated data, and snRNA-seq data intron retention fraction was approximated at 0.4

Supplemental Table 2: Overall F1 accuracy scores across software packages.

F1 scores for all benchmarked software on all simulated data.

Supplemental Table 3: F1 Accuracy scores as a function of allowed multimapper alignments.

F1 scores for TEsingle on simulated data allowed 1, 10, 100, or 1000 mapping locations per read.

Supplemental Table 4: Benchmarked accuracy on genes

Accuracy breakdowns for gene annotations on simulated data for all benchmarked software.

**Supplemental Table 5: Benchmarked accuracy on TEs by subfamily** Accuracy breakdowns for TE annotations by subfamily on simulated data for all benchmarked software.

Supplemental Table 6: Benchmarked accuracy on TEs by locus

Accuracy breakdowns for TE annotations by individual locus on simulated data for all benchmarked software.

Supplemental Table 7: Martirosyan donor tissue metadata and single-nuclei RNA-seq recovery counts.

Donor tissue metadata and single-nuclei RNA-seq recovery counts used from the Martirosyan analysis. Metadata of donor tissues includes donor diagnosis, age, sex, and the total numbers of recovered nuclei across broad cellular populations and subpopulations after quality control.

Supplemental Table 8: List of DE genes and TEs between broad cell type populations.

List of DE genes and TEs found between broad cell type populations using a logistic regression framework. Significantly DE genes and TE loci were identified with threshold of p_val_adj < 0.05.

Supplemental Table 9: List of cellular subpopulation cluster markers identified through Leiden subpopulation re-clustering.

List of cellular subpopulation cluster markers. **A:** Neuronal subpopulation cluster markers found with Seurat’s FindAllMarkers. Each subpopulation is denoted with a header and genes are listed along with the found p-value (P_val), average log2 fold change (Avg_log2FC), the percentage of cells expressing the gene within the denoted subpopulation (Pct_1), the percentage of cells express the gene in all other subpopulations (Pct_2), the adjusted p-value (P_val_adj), and direction of expression of the gene (Regulation). **B:** Astrocytic state cluster markers. **C:** Microglial state cluster markers. **D:** Oligodendrocyte subpopulation cluster markers.

Supplemental Table 10: List of DE genes and TEs of PD vs control neuronal subpopulations.

List of DE genes and TEs of PD vs control neuronal subpopulation with Seurat’s FindMarkers. DE analysis was performed to find DE genes and TEs between PD and control cells from the same neuronal subpopulations. A category tag was created to identify clinical diagnosis and subpopulation cluster. Then a loop was used to perform the differential expression analysis using a logistic regression framework. Significantly DE genes and TE loci were identified with threshold of p_val_adj < 0.05. **A:** DopaN DE genes and TEs between PD and control. **B:** Inhibitory neurons DE genes and TEs between PD and control. **C:** Excitatory neurons DE genes and TEs between PD and control. **D:** Other neurons DE genes and TEs between PD and control.

Supplemental Table 11: List of DE genes and TEs of PD vs control astrocytic substates.

List of DE genes and TEs of PD vs control astrocytic substates with Seurat’s FindMarkers. DE analysis was performed to find DE genes and TEs between PD and control cells from the same astrocytic substate. A category tag was created to identify clinical diagnosis and subpopulation cluster. Then a loop was used to perform the differential expression analysis using a logistic regression framework. Significantly DE genes and TE loci were identified with threshold of p_val_adj < 0.05. **A:** Homeostatic astrocytes DE genes and TEs between PD and control. **B:** Reactive astrocytes DE genes and TEs between PD and control. **C:** Disease associated astrocytes DE genes and TEs between PD and control. **D:** Inflammatory astrocytes DE genes and TEs between PD and control. **E:** Metallothionein astrocytes DE genes and TEs between PD and control.

Supplemental Table 12: List of DE genes and TEs of PD vs control microglial substates.

List of DE genes and TEs of PD vs control microglial substates with Seurat’s FindMarkers. DE analysis was performed to find DE genes and TEs between PD and control cells from the same microglial substate. A category tag was created to identify clinical diagnosis and subpopulation cluster. Then a loop was used to perform the differential expression analysis using a logistic regression framework. Significantly DE genes and TE loci were identified with threshold of p_val_adj < 0.05. **A:** Homeostatic microglia (HM) DE genes and TEs between PD and control. **B:** Cytokine responding microglia (CRM) DE genes and TEs between PD and control. **C:** Interferon responding microglia (IRM) DE genes and TEs between PD and control. **D:** Antigen presenting microglia (APM) DE genes and TEs between PD and control. **E:** Disease associated microglia (DAM) DE genes and TEs between PD and control.

Supplemental Table 13: List of DE genes and TEs of PD vs control oligodendrocyte subpopulations.

List of DE genes and TEs of PD vs control oligodendrocyte (ODCs) subpopulations with Seurat’s FindMarkers. DE analysis was performed to find DE genes and TEs between PD and control cells from the same oligodendrocyte subpopulations. A category tag was created to identify clinical diagnosis and subpopulation cluster. Then a loop was used to perform the differential expression analysis using a logistic regression framework.

Significantly DE genes and TE loci were identified with threshold of p_val_adj < 0.05. **A:** Resting ODCs DE genes and TEs between PD and control. **B:** Myelinating ODCs DE genes and TEs between PD and control. **C:** Immature ODCs DE genes and TEs between PD and control. **D:** Stress associated ODCs DE genes and TEs between PD and control. **E:** Ox stress associated ODCs DE genes and TEs between PD and control.

Supplemental Table 14: Key Resource Table

Data and code used and generated in this study.

## Resource Availability

### Data and code availability

The data, code, protocols, and key lab materials used and generated in this study are listed in a Key Resources Table alongside their persistent identifiers in **Supplemental Table 14**.

An earlier version of this manuscript was posted to **[preprint server]** on **[date] at [DOI].**

The following published single-nuclei datasets were obtained from Gene Expression Omnibus (accessions: GSE140231, GSE243639). FOUNDIN-PD single-cell datasets were obtained from the FOUNDIN-PD group (https://www.foundinpd.org/). Simulated datasets used for benchmarking has been deposited in Zenodo (DOI: 10.5281/zenodo.18261667, https://doi.org/10.5281/zenodo.18261667).

All original code used for this study are available at GitHub (https://github.com/mhammell-laboratory/TEsingle, https://github.com/mhammell-laboratory/TEsingle_benchmarking_scripts, and https://github.com/mhammell-laboratory/TEsingle_Martirosyan_analysis_code), and have been deposited in Zenodo (**Supplemental Table 14**).

## Acknowledgements

We wish to thank Agnete Kirkeby and her team for helpful discussions. This research was funded by Aligning Science Across Parkinson’s (https://ror.org/03zj4c476) (Grant #: ASAP-000520, ASAP-024296 and ASAP-025170) through the Michael J. Fox Foundation for Parkinson’s Research (MJFF) (https://ror.org/03arq3225). Funding was also provided in part by the CZI Neurodegeneration Challenge Network. For the purpose of open access, the author has applied a CC BY public copyright license to all Author Accepted Manuscripts arising from this submission.

## Author Information

These authors contributed equally: Talitha Forcier, Esther Cheng.

## Contributions

MGH conceived, funded and supervised the study. TF designed and implemented the methodology. TF, OT, and CW designed and performed the benchmarking analysis. EC and OT participated in data collection and computational analysis. TF and EC designed and generated all figures. All authors contributed to manuscript writing and revising. All authors read and approved the final version of the manuscript.

---

**Supplemental Figure 1:**
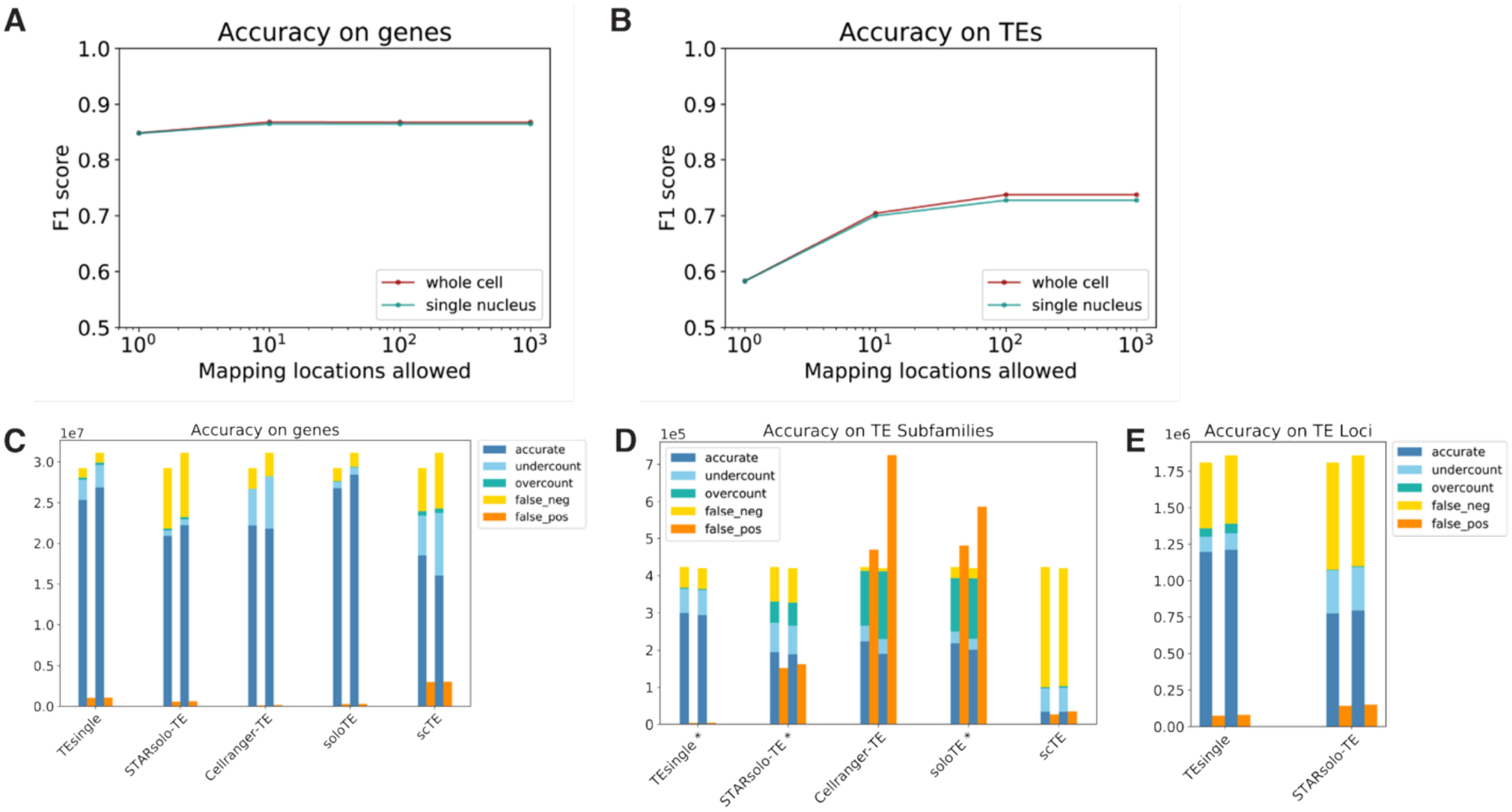
Further accuracy measurements of benchmarking results on simulated data. **A:** TEsingle F1 scores for gene expression evaluation when reads are allowed at most 1 mapping location, 10 mapping locations, 100 mapping locations, and 1000 mapping locations for simulated whole-cell and snRNAseq data. **B:** TEsingle F1 scores for TE expression evaluation when reads are allowed at most 1 mapping location, 10 mapping locations, 100 mapping locations, and 1000 mapping locations for simulated whole-cell and snRNAseq data. **C-E:** Raw counts for the number of annotations within each cell evaluated accurately (measured expression within 15% of simulated expression), overestimated (measured expression greater than 115% of simulated expression), underestimated (measured expression less than 85% of simulated expression), false negative (measured to have zero expression when nonzero expression simulated), and false positive (measured to have nonzero expression when zero expression simulated) for each benchmarked software on both whole-cell and snRNAseq simulated data (left and right pairs, respective of each software’s set of bar plots’). These values were then used to calculate F1 scores plotted in the main body of the paper. **C:** Results for gene annotations. **D:** subfamily-level results for TE annotations. **E:** locus-specific results for TE annotations.

**Supplemental Figure 2:**
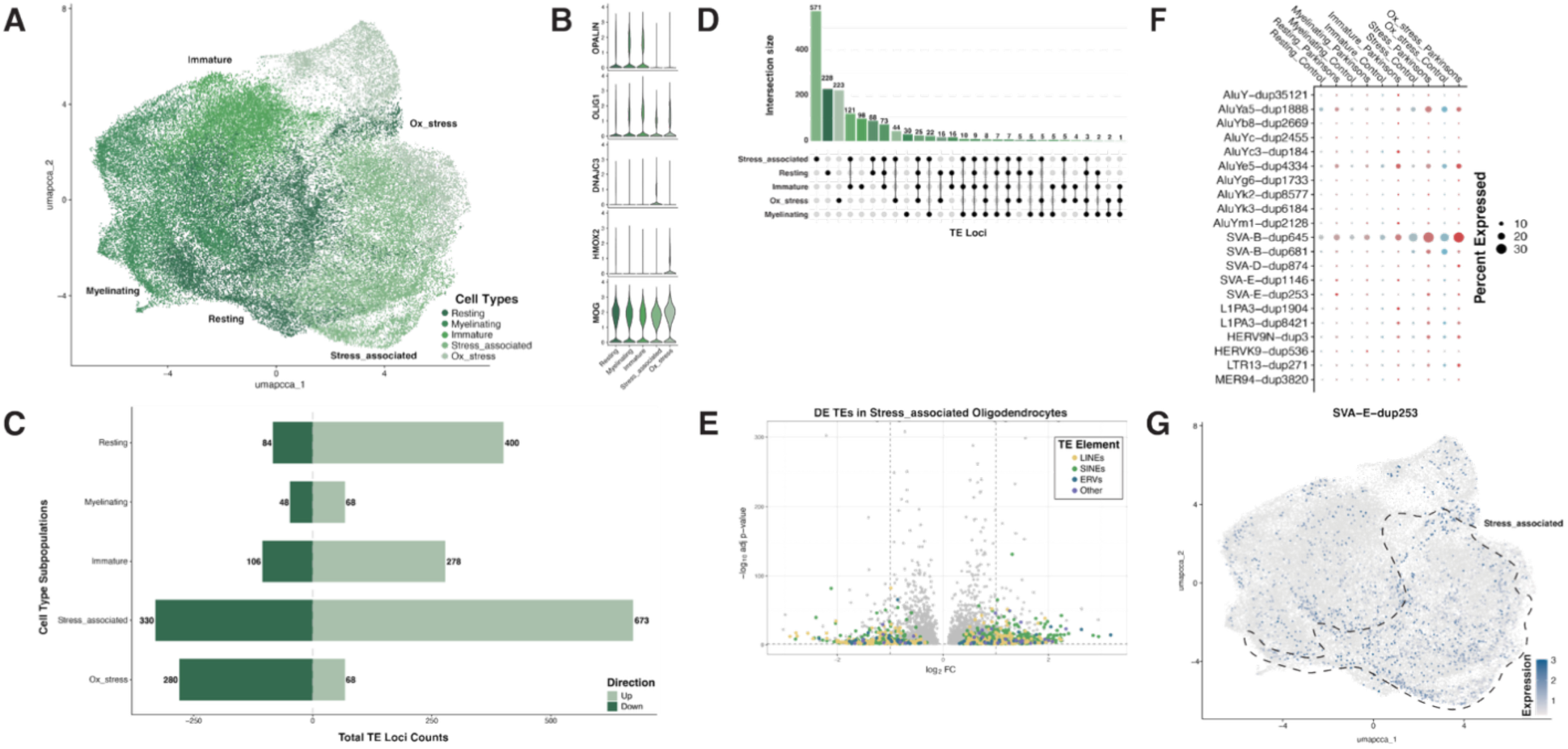
PD oligodendrocytes show an elevated stress response state marked by an enrichment of young and human-specific TE loci. **A:** UMAP based Leiden clustering of all oligodendrocytes in the Martirosyan dataset showing the identification of five ODC subpopulations: resting, myelinating, immature, stress-associated, and Ox-stress associated ODCs. **B:** Violin plots displaying the marker genes that are enriched within each ODC subpopulation. **C:** Horizontal bar plot counting the total number of DE TE loci within each PD-derived ODC subpopulation after differential expression analysis to the matched control ODC subpopulation. **D:** Upset plot showing the total counts of DE TE loci within each PD-derived ODC subpopulation or shared between ODC subpopulations. Counts includes both up and downregulated DE TE loci. **E:** Volcano plot of fold change versus negative log10 adjusted p-values displaying all significantly DE TE loci (colored dots) against all significantly DE genes (gray dots) within PD-derived stress associated ODCs. Each colored point is a specific TE locus that is colored based on the TE class they belong to, as noted in the legend. **F:** Dot plot showing the percent expression of selected younger and human-specific TE loci between PD and control ODC subpopulations. The size of the dot represents the percentage of cells that express the TE locus, and the color encodes the diagnosis of PD (red) or control tissue (blue). **G:** Feature plot showcasing the SVA-E-dup253 TE locus that is specifically enriched within stress associated ODCs. Colored dots represent single cells on the oligodendrocyte UMAP presented in **Supplemental** Figure 2A, where the darker the colored cells, the higher the expression value of the SVA-E-dup253 TE locus within that cell.

